# Phosphorylation of the canonical histone H2A marks foci of damaged DNA in malaria parasites

**DOI:** 10.1101/2020.11.06.372391

**Authors:** Manish Goyal, Adina Heinberg, Vera Mitesser, Sofia Kandelis-Shalev, Brajesh Kumar Singh, Ron Dzikowski

## Abstract

*Plasmodium falciparum* parasites proliferate within circulating red blood cells and are responsible for the deadliest form of human malaria. These parasites are exposed to numerous intrinsic and external sources that could cause DNA damage, therefore, they have evolved efficient mechanisms to protect their genome integrity and allow them to proliferate in such conditions. In higher eukaryotes, double strand breaks rapidly lead to phosphorylation of the core histone variant H2A.X which marks the site of damaged DNA. We show that in *P. falciparum* that lacks the H2A.X variant, the canonical PfH2A is phosphorylated on serine 121 upon exposure to sources of DNA damage in a dose dependent manner. We further demonstrate that phosphorylated PfH2A is recruited to foci of damaged chromatin shortly after exposure to sources of damage, while the non-phosphorylated PfH2A remains spread throughout the nucleoplasm. In addition, we found that PfH2A phosphorylation is dynamic and as the parasite repairs its DNA over time, this phosphorylation is removed. We also demonstrate that these phosphorylation dynamics could be used to establish a novel and direct DNA repair assay in *P. falciparum*.

**Importance:** *Plasmodium falciparum* is the deadliest human parasite that causes malaria when it reaches the blood stream and begins proliferating inside red blood cells where the parasites are particularly prone to DNA damage. The molecular mechanisms that allow these pathogens to maintain their genome integrity under such condition are also the driving force for acquiring genome plasticity that enable them to create antigenic variation and become resistant to essentially all available drugs. However, mechanisms of DNA damage response and repair have not been extensively studied in these parasites. The paper addresses our recent discovery, that *P. falciparum* that lacks the histone variant H2A.X, phosphorylates its canonical core histone PfH2A in response to exposure to DNA damage. The process of DNA repair in Plasmodium was mostly studied indirectly. Our findings enabled us to establish a direct DNA repair assay for *P. falciparum* similar to assays that are widely used in model organisms.

## Introduction

*Plasmodium falciparum* is the protozoan parasite responsible for the deadliest form of human malaria. This parasite is estimated to infect 200-300 million people worldwide each year, resulting in approximately half a million deaths, primarily of young children [1]. *P. falciparum* replicates within the circulating red blood cells of an infected individual, and its virulence is attributed to its ability to modify the erythrocyte surface and to evade the host immune attack. During their intra-erythrocytic development, *Plasmodium* parasites replicate their haploid genomes multiple times through consecutive mitosis cycles called schizogony, which makes them particularly prone to errors during DNA replication. In addition, blood stage parasites that live in a highly oxygenated environment produce potent DNA damaging agents while digesting hemoglobin and are exposed to oxidative substances released from immune cells [2].

Therefore, *Plasmodium* parasites that are exposed to numerous sources that can damage their DNA must have evolved efficient mechanisms to protect their genome integrity. Orthologues to many of the proteins involved in the DNA damage response (DDR) are encoded in *P. falciparum* genome [2] including those involved in homologous recombination (HR), microhomology-mediated end joining (MMEJ) [2] and mismatch repair machineries [3]. However, these mechanisms have not been extensively studied in these parasites. It appears that malaria parasites utilize both HR and an alternative end joining pathway to maintain their genome integrity [4]. Thus, in the absence of a homologous sequence in their haploid genome that can serve as a template for HR, blood stage parasites primarily repair double strand brakes (DSB) using the alternative microhomology-mediated end joining mechanism (MMEJ) [4, 5].

In mammals, a single double-strand break of the DNA triggers the DDR that rapidly leads to extensive ATM-kinase-dependent phosphorylation of the core histone isoform H2A.X to form a phospho-H2A.X (γ-H2A.X), which marks the site of damaged DNA [6]. However, the *P. falciparum* genome lacks an orthologue of the H2A.X variant, and it only encodes two H2A variants, the canonical PfH2A (PF3D7_0617800) and PfH2A.Z (PF3D7_0320900), which was shown to be associated with subset of active promoters [7]. Previous histone phosphorylation analysis suggested that PfH2A could be phosphorylated on serine 121 [8]. We were interested to determine whether in *P. falciparum* phosphorylation of PfH2A might be correlated with DNA damage. We show that these parasites phosphorylate the canonical PfH2A on serine 121 in response to DNA damage and that the phosphorylated PfH2A is recruited to the damaged foci. In addition, the ability to specifically detect the dynamics of this phosphorylation using an anti γ-H2A.X antibody provides a useful marker for studying DNA damage mechanisms which allowed us to establish a direct DNA repair assay in *P. falciparum*.

## Results

In model systems, phosphorylation of H2A.X is elevated following exposure to DNA damaging agents and is commonly used as a marker for double strand breaks [6]. The phosphorylation of serine 139 found on a conserved SQ (Serine-Glutamine) motif of mammalian H2A.X serves as a differential epitope for detection of the phosphorylated form known as γ-H2A.X. *Plasmodium* parasites do not contain a gene encoding the H2A.X variant in their genome, but instead they express the canonical H2A and the H2A.Z variants (PF3D7_0617800 and PF3D7_0320900 respectively). In the absence of a good marker for DNA damage in *Plasmodium,* we were interested to test whether the antibody that recognizes γ-H2A.X in mammals could be used as a marker for DNA damage in *P. falciparum*. As a first step we aligned the two plasmodium H2A variants with human H2A.X and noted that only the canonical PfH2A has a long C-terminal tail containing the SQ motif, which is conserved among *Plasmodium* species, while no SQ motif is found in PfH2A.Z (Fig. S1). This SQ motif is conserved among the canonical H2A of several protozoan parasites such as *P. falciparum*, *Giardia lamblia* and *Trichomonas vaginalis* that lack H2A.X orthologues. Similarly, the budding yeast *Saccharomyces cerevisiae* is lacking the H2A.X orthologue, and instead its canonical H2A was found to be phosphorylated on serine 129 (SQ motif). This phosphorylation is detected by the anti-γ-H2A.X antibody, and thus, the phosphorylated form of *S. cerevisiae* H2A is often referred to as γ-H2A.X [9]. Interestingly, contrary to *Plasmodium spp*., the apicomplexan parasite *Toxoplasma gondii* has an H2A.X variant in addition to its canonical H2A (Fig. 1A). As expected, *in silico* structural prediction of PfH2A suggests that the SQ motif is found in its C’ terminal tail (Fig. 1B) and that this motif is likely to be an ATM kinase phosphorylation site (Fig. 1C), similar to the serine 139 of the mammalian H2A.X, which is the major residue phosphorylated in response to DNA damage by the ATM kinase [6].

**Figure 1.**
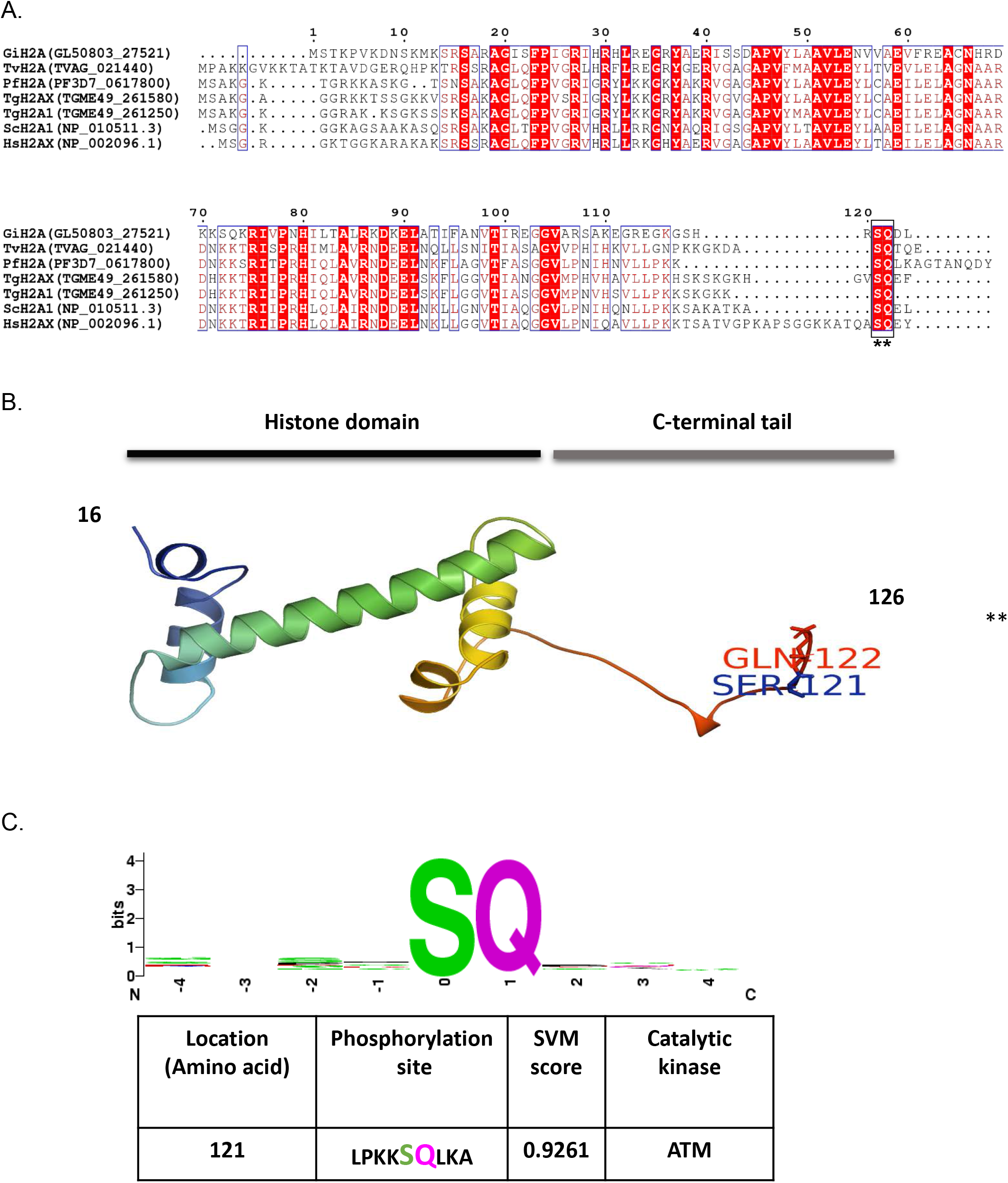
*In silico* analyses of putative ATM kinase-specific phosphorylation site (conserved SQ-motif) in PfH2A. **(A)**. Multiple sequence alignment of amino acid sequences of PfH2A and some of H2A variants from human, budding yeast, and protozoan parasites using ESPript 3. Similar and identical amino acids are boxed and marked with a red background. The conserved C’-terminal SQ motif is underlined with asterisks. The species names and corresponding uniport accession number are as follows: PfH2A; *P. falciparum* histone H2A, HsH2AX; *Homo sapiens* Histone H2AX, ScH2A1: *Saccharomyces cerevisiae* Histone H2A1, TgH2A1; *Toxoplasma gondii* Histone H2A1, TgH2AX; *Toxoplasma gondii* Histone H2A.X, TvH2A; *Trichomonas vaginalis* Histone H2A, and GiH2A; *Giardia intestinalis* Histone H2A. **(B).** 3D-Homology model of PfH2A (developed using structure homology-modeling server SWISS-MODEL) showing core histone domain and extended C-terminal tail contains the conserved S^121^Q^122^ motif. **(C)**. *In silico* prediction of ATM kinase-specific phosphorylation sites in PfH2A (using KinasePhos, version 2.0). The sequence-based amino acid coupling-pattern analysis and solvent accessibility of PfH2A suggest Serine 121 as the most prominent ATM kinase specific phosphorylation site.

To test if PfH2A is indeed phosphorylated in response to exposure of the parasite to a source of DNA damage, we exposed tightly synchronized ring stage NF54 parasites to X-ray irradiation. We chose to irradiate early stage parasites that are not replicating their DNA and do not have haemozoin, and therefore the detected DNA damage should be mostly due to the exogenous source. We used TUNEL assays as direct evidence that exposure of the parasite to 6000 Rad caused DNA damage, which was detected in most parasite’s nuclei (Fig 2A). We then used the γ-H2A.X antibody for immuno-fluorescence assays (IFA) and were able to detect strong signals within the parasites’ nuclei after exposure to X-ray irradiation (Fig 2B). This observation was further confirmed using γ-H2A.X antibody on proteins extracted from parasites exposed to increasing levels of irradiation which showed a corresponding elevation in the levels of γ-PfH2A (Fig. 2C, left panel). Similarly, exposing the parasites to H_2_O_2_, another source of DNA damage, caused an increase in the levels of γ-PfH2A recognition (Fig. 2C, right panel). To ensure that the anti-γ-H2A.X antibody specifically recognized the phosphorylated form of PfH2A and did not cross-react with the non-phosphorylated form, we incubated the extracted proteins with calf intestine phosphatase (CIP) that removes phosphate residues. The CIP treatment specifically abolished immunoblot detection using the anti-γ-H2A.X antibody while the non-phosphorylated PfH2A was detected at similar levels in parasites exposed to increasing X-ray levels (Fig 2D). In addition, when we initially probed with the anti-γ-H2A.X antibody after irradiation, we observed increasing levels of phosphorylation, however, when the blot was stripped, treated with CIP and re-probed, the anti-γ-H2A.X antibody signal disappeared while detection of the canonical PfH2A was unchanged (Fig. 2E). Altogether, these data suggest that PfH2A is phosphorylated in response to DNA damage and that the anti-γ-H2A.X antibody is specific to the phosphorylated form of PfH2A.

**Figure 2.**
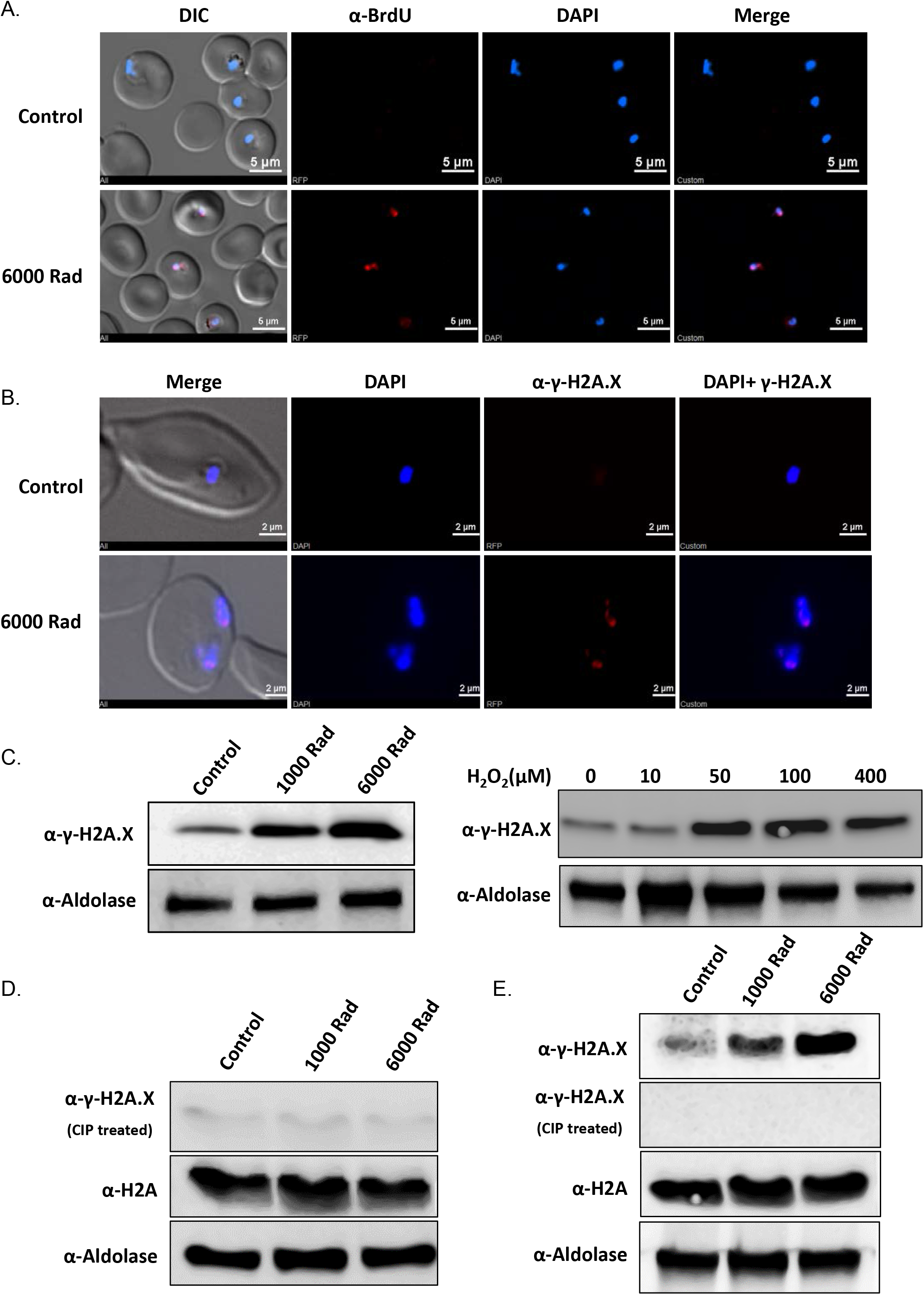
DNA damage in *P. falciparum* causes histone H2A phosphorylation in a dose-dependent manner. **(A).** DNA fragmentation imaging by TUNEL assay of RBCs infected with NF54 *P. falciparum* parasites exposed to X-ray irradiation (6000 rad) showing nuclear foci of damaged DNA. **(B)**. Immunofluorescence analysis of X-ray irradiated parasites (6000 rad), using anti-γ-H2A.X(S^P^Q) antibody, shows foci of phosphorylation signal in the nucleus. **(C).** Western blot analysis, using anti-γ-H2A.X(S^P^Q) antibody, of protein extracts from parasites exposed to increasing levels of X-ray irradiations (left) and from parasites treated with increasing concentrations of H_2_O_2_ (right). **(D-E)**. The anti-γ-H2A.X (S^P^Q) antibody specifically recognizes the phosphorylated PfH2A and does not cross-react with non-phosphorylated PfH2A. Protein from parasites, which were irradiated with increasing doses of X-ray radiation (control (no irradiation), 1000, and 6000 Rad respectively) were subjected to WB analysis using either anti-γ-H2A.X (S^P^Q) antibody or anti-H2A antibody. The membrane was either incubated with calf intestine phosphatase (CIP) before incubation with the antibodies (D) or incubated with the antibodies, stripped, treated with (CIP) and re-incubated with the antibodies (E). anti-Aldolase antibody was used as a loading control. The anti-γ-H2A.X (S^P^Q) antibody detected increasing levels of protein associated with the increasing levels of irradiation only without CIP treatments while the anti-H2A antibody detected constant protein levels even after CIP treatment.

To further confirm that the phosphorylation detected by the anti-γ-H2A.X antibody, following parasite’s exposure to DNA damage, is indeed phosphorylation of the canonical PfH2A, we extracted and purified histones from parasites that were either exposed or not exposed to X-ray irradiation. Immunoblot analysis of the total histone extract shows an increase in the level of the phosphorylated form of PfH2A following irradiation while the total levels of PfH2A are similar (Fig. 3A & B). We further exposed parasites to X-ray irradiation and performed immuno-precipitation (IP) of total PfH2A using an anti-H2A antibody. The IP fractions were subjected to immunoblot with the anti-γ-H2A.X antibody, which demonstrated significant enrichment of the phosphorylated form of PfH2A in the elution (Fig. 3C). This fraction was subjected to trypsin digestion followed by mass spectrometry analysis, which identified phosphorylation on serine 121 of PfH2A (Fig. 3D).

**Figure 3.**
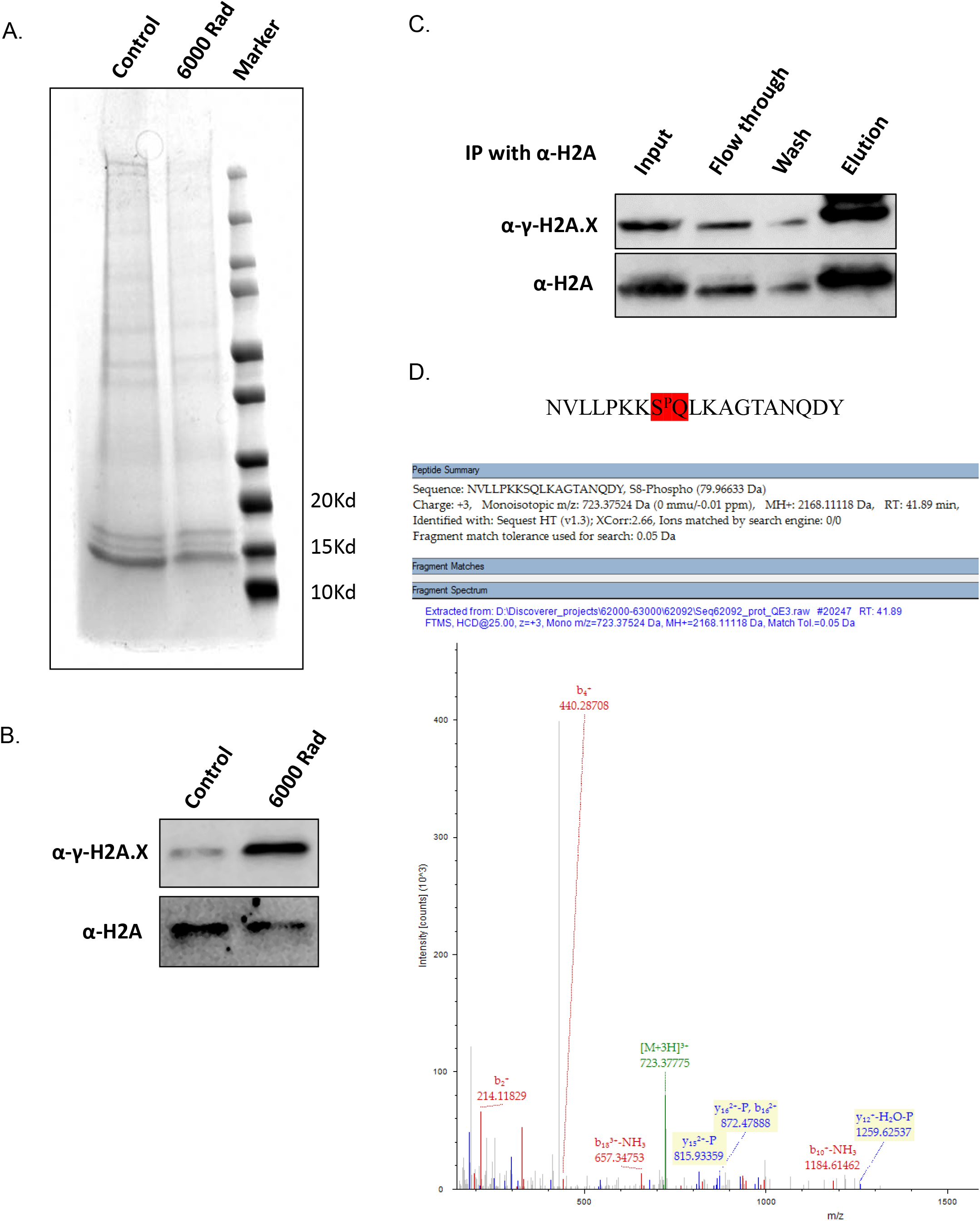
Histone extraction followed by Mass spectrometry shows that Serine 121 of PfH2A is phosphorylated upon exposure to X-ray irradiation. **(A)**. SDS page analysis of histone extraction and purification from X-ray irradiated (6000 rad) and untreated parasites. **(B).** Western blot analysis of total histone extracted from untreated and X-ray treated (6000 Rad) parasite using anti-γ-histone H2AX (S^P^Q) and anti-PfH2A antibodies. **(C)**. X-ray irradiated parasites (6000 Rad) were subjected to Immunoprecipitation (IP), using anti-H2A antibody, followed with WB analysis using both anti-γ-H2A.X (S^P^Q) and anti-H2A antibodies. **(D).** Trypsin digestion followed by mass spectrometry analysis identified phosphorylation of Serine 121 of PfH2A in the irradiated parasites.

The specificity of the anti γ-H2A.X antibody to the phosphorylated form of PfH2A allowed us to image its nuclear distribution compared with the non-phosphorylated PfH2A. Immuno fluorescence assay using anti H2A and anti γ-H2A.X antibodies indicated that while the canonical PfH2A is spread throughout the nucleoplasm, its phosphorylated form is found at distinct foci (Fig. 4A). To further validate this observation, we performed super resolution STORM imaging that enabled us to image the nuclear distribution of the two forms of PfH2A in detail at the nanoscale level. This analysis clearly demonstrates the differential distribution of the two PfH2A forms in the nucleoplasm. The non-phosphorylated PfH2A is indeed spread throughout the nucleoplasm while the phosphorylated form is much less abundant, and is found at distinct nuclear foci (Fig. 4B).

**Figure 4.**
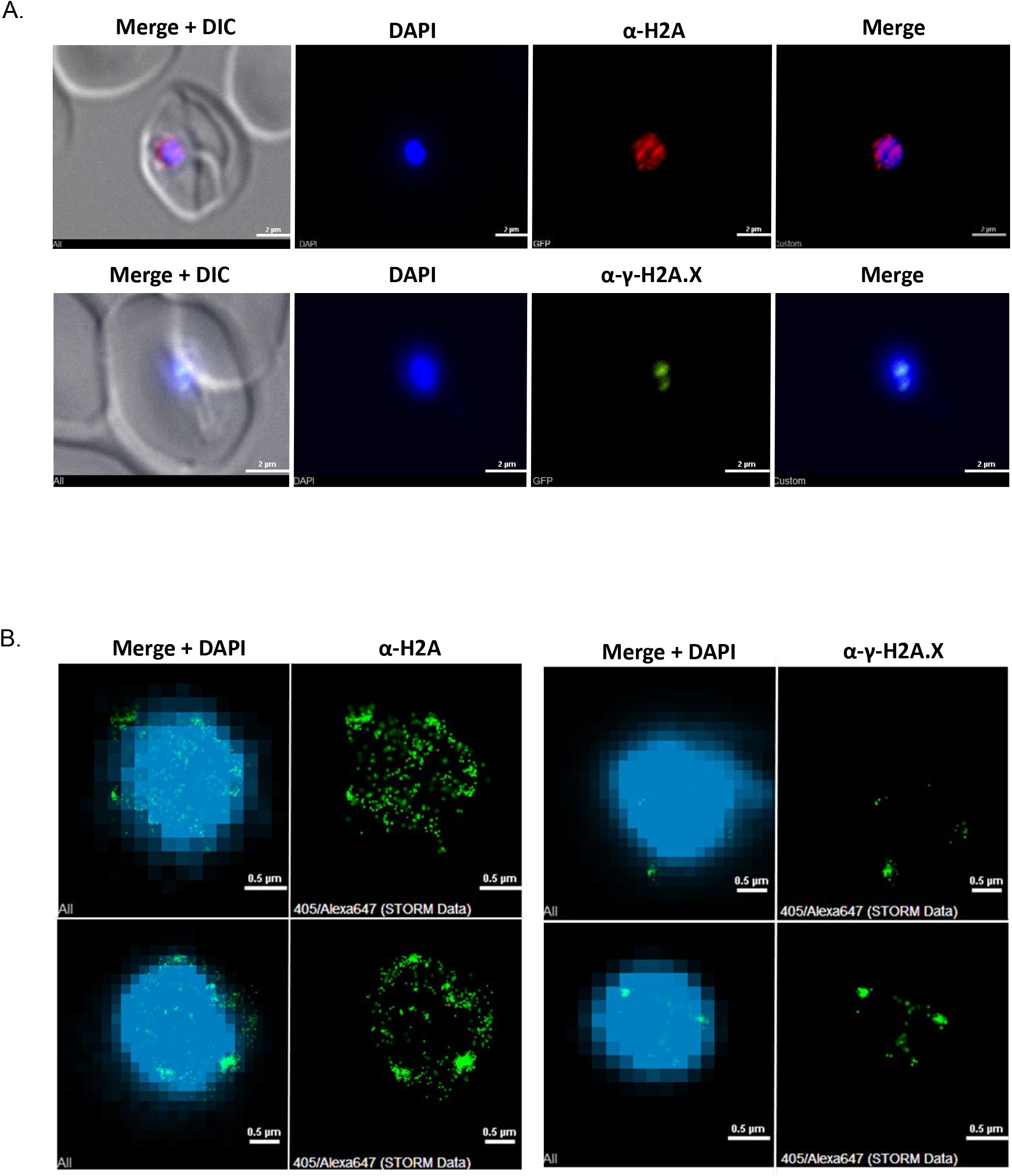
Phosphorylated PfH2A is located at distinct nuclear foci while the non-phosphorylated PfH2A is spread throughout the nucleoplasm. **(A)**. Immunofluorescence (red, α-γ-H2A; green, α-γ-H2A.X; blue, DAPI. Scale bar 2 μm) and **(B)** Super resolution STORM imaging of PfH2A and phosphorylated PfH2A (green, Alexa 647 staining each of the PfH2A isoforms; blue, YOYO1 staining of DNA at low resolution for orientation. Scale bar 0.5 μm) in *P. falciparum* nuclei following X-ray irradiation (6000 Rad).

Thus far, the process of DNA repair in Plasmodium was mostly studied indirectly by measuring the recovery of parasites in culture after exposure to a source of DNA damage [10]. In addition, repair mechanisms were studied directly by creating a transgenic inducible DSB system by integrating an *I SceI* cleavage site into the *P. falciparum* genome and sequencing of the repaired locus after induction of the *I SceI* endonuclease [4]. Our data strongly suggest that phosphorylation of PfH2A could be used as a specific and immediate marker for damaged DNA in *P. falciparum.* Therefore, we were interested to examine the dynamics of this phosphorylation over time after exposure to X-ray irradiation, hypothesizing that it could be exploited to establish a direct DNA repair assay for *P. falciparum* similar to assays that are widely used in model organisms. We exposed parasite cultures to different levels of X-ray irradiation and measured the levels of PfH2A phosphorylation over time. We observed that the levels of PfH2A phosphorylation, which had increased immediately after irradiation, decreased already 3 hours after irradiation to levels that are similar to those prior to irradiation (Fig. 5). These data imply that during this period of time, the parasites were able to repair their damaged DNA, and thus, these dynamics could be used as a valuable tool to study DNA repair in malaria parasites.

**Figure 5.**
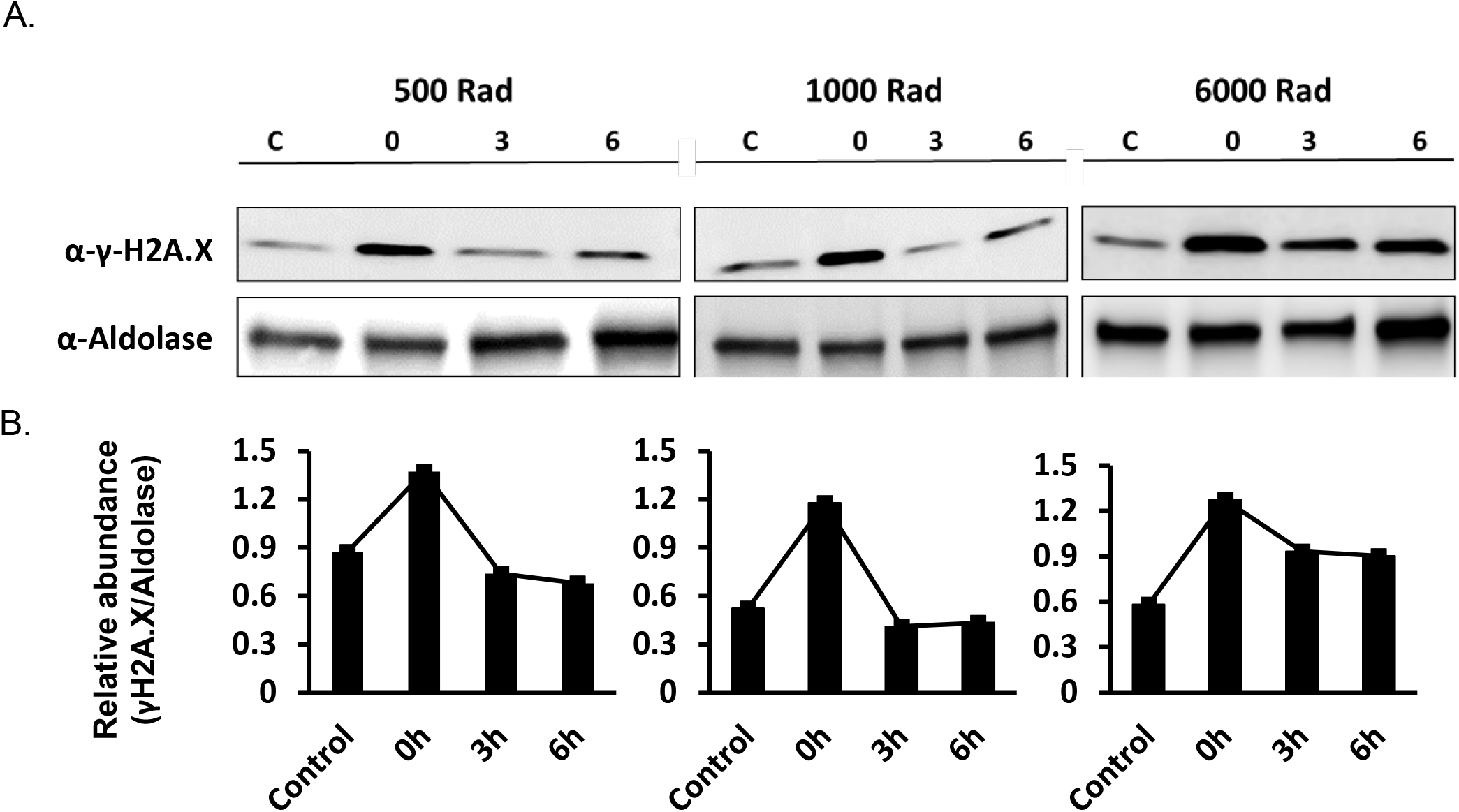
DNA damage and repair assay in *P. falciparum* iRBCs. (**A).** Parasites were treated with different doses of X-ray radiation (i.e. 500, 1000 and 6000 Rad) and put back in culture (3h and 6h) to allow them to repair their damaged DNA. Protein extracts from these parasites were then used for WB analysis with anti-γ-H2A.X (S^P^Q) antibody and anti-aldolase antibody as a loading control. The ability of the parasites to repair their damaged DNA is demonstrated by the rapid reduction in the levels of phosphorylated PfH2A found 3h after irradiation. **(B).** Semi-quantitative densitometry analysis of the WB presented in (A). Quantification of the changes in the ratio between the signal detected by the anti-γ-H2A.X and anti-aldolase antibody before irradiation, immediately after irradiation, 3h and 6h after irradiation. The density of each band is presented as a proportion of the total signal obtained. C, parasites that were not exposed to irradiation and used as control.

## Discussion

In any living organism, the ability to repair damaged DNA is key for maintaining genome integrity. This DNA damage repair (DDR) machinery should be extremely efficient in organisms such as *Plasmodium* parasites that are continuously exposed to numerous intrinsic and exogenous sources that may damage their DNA. In addition to its crucial role for the parasite’s basic biological functions under these conditions, efficient DDR machinery contributes to the parasite’s ability to expand its antigenic repertoire and to maintain mutations that enable it to resist drug treatment. However, although many regulators of DDR were identified encoded in the *Plasmodium* genome, the mechanisms for DDR in these parasites remained understudied and poorly understood. A major obstacle for advancing our knowledge on DDR machinery in *Plasmodium* is the lack of good molecular markers for damaged DNA and the inability to perform an accurate assay that directly measures the kinetics of DNA repair. Here we show that exposure of *P. falciparum* parasites to X-ray irradiation and H_2_O_2_, which cause double-strand breaks (DSB), leads to phosphorylation of the canonical PfH2A in a dose dependent manner. We found that although *Plasmodium* has no H2A.X variant, the canonical PfH2A is phosphorylated on the SQ motif found in its C’-terminal-tail and that the phosphorylated PfH2A could be differentiated from the non-phosphorylated form of this core histone protein. Since PfH2A is phosphorylated in a dose-dependent manner, the quantitative measurement of phosphorylated PfH2A can act as a sensitive molecular marker for DNA damage in *P. falciparum*. Most of the approaches employed to date to study DNA damage in *Plasmodium* rely on measuring the relative instability of DNA damage products under alkaline conditions (comet assay) [11, 12] or on the relative expression of DNA damage and repair genes (qRT-PCR) [11, 13, 14]. In addition, thus far, the ability of *P. falciparum* parasites to repair DNA damage was estimated by the rate of recovery of parasite populations exposed to DNA damaging agents i.e. the time it takes for these populations to reach approximately 5% parasitemia (usually 10-20 days) [10]. Any delay in recovery was then interpreted as a malfunction of the repair machinery, which is of course indirect evidence reflected two weeks after the actual repair has happened. In this manner, the analysis of PfH2A phosphorylation kinetics can fulfill the need for a direct, simple, sensitive/quantitative and reproducible way of measuring DNA damage and repair kinetics in Plasmodium in a time scale of minutes to hours, which better represents the velocity of the repair machinery.

In higher organisms, phosphorylation of histone variant H2A.X is a highly specific and sensitive molecular marker for monitoring DNA damage and repair [15, 16]. However, in some organisms, other histone H2A variants undergo phosphorylation in response to exposure to DNA damage. For example, in *Drosophila melanogaster* H2A.Z is phosphorylated in response to DNA damage instead of H2A.X [17], and in the budding yeast *Saccharomyces cerevisiae* that do not encode an H2A.X variant, the canonical H2A is phosphorylated at the serine found near its C-terminus at an SQ motif [18], similar to what we report here in *P. falciparum*. This also appears to be the case in protozoan species where histone H2A.X is either missing or replaced by other histone variants. A marked example is in *Trypanosoma brucei* and other trypanosomatids as well, in which histone H2A undergoes phosphorylation at a threonine residue (Thr 130) instead of serine in response to DNA damage [19]. Interestingly, in the apicomplexan parasite *Toxoplasma gondii*, which does contain an H2A.X variant, the canonical H2A (also named H2A1) was also proposed to be phosphorylated at a C-terminal SQ motif as a response to DSBs [20]. This may suggest the possibility of functional redundancy among these variants that could be exploited through evolution for functional replacement by the canonical H2A when the H2A.X variant is missing. This is somehow supported by the high level of conservation of the SQ motif in the canonical H2A of other protozoan and in particular in other *Plasmodium* species that face similar exposure to sources of DNA damage such as *P. falciparum.* Interestingly, lower eukaryotes prefer high fidelity HR as the mechanisms to repair DSB while higher eukaryotes prefer NHEJ [21]. We noted that both *P. falciparum* and *S. cerevisiae* that phosphorylate their canonical H2A in response to DSB use HR for repair, while *T. gondii* that encode and phosphorylate TgH2A.X variant use mainly NHEJ similar to higher eukaryotes [21, 22]. One can speculate whether the preference for NHEJ might have evolved in association with the H2A.X variant.

In higher eukaryotes, histone H2A.X is known to be phosphorylated by members of phosphatidylinositol 3-kinase family (PI3K) namely Ataxia Telangiectasia Mutated (ATM) kinase, ATM Rad-3-related kinase (ATR), and DNA-dependent protein kinase (DNA-PK) [23]. However, to the best of our knowledge only one PI3K was identified in Plasmodium [24] while other members of this family are not well characterized. Interestingly, in the apicomplexan parasite *T. gondii*, an ATM kinase orthologue (TGME49_248530) was proposed to be involved in TgH2A.X phosphorylation [25]. Incubation of cultured parasites with a known ATM kinase inhibitor (KU-55933), which was tested as a potential anti *T. gondii* agent, caused cell cycle arrest and was able to inhibit phosphorylation of TgH2A.X [26]. The single PI3K which was previously identified in *P. falciparum* (PF3D7_0515300), shows some sequence conservation with *T. gondii* (TGME49_248530) and Human (AAB65827) ATM kinases (Fig. S2), and incubation of parasites with KU-55933 was able to reduce the levels of PfH2A phosphorylation after irradiation (Fig S3). However, this *P. falciparum* PI3K kinase (PF3D7_0515300), which was shown to be important for hemoglobin digestion was localized to the parasite Plasma Membrane (PM), Parasitophorous Vacuole Membrane (PVM) and the Food Vacuole (FV) but did not appear to be localized to the nucleus [24]. Thus, the plasmodium ATM kinase homologue that phosphorylates PfH2A is yet to be identified.

Overall, the identification of PfH2A phosphorylation as a marker for DNA damage and the ability to quantify and time the appearance and disappearance of this marker in response to exposure to sources of DNA damage, opens new possibilities for understanding the mechanisms of DNA damage and repair that contribute to the persistence and pathogenicity of these important pathogens.

## Materials and Methods

### Parasite culture

All experiments were conducted on the human malaria NF54 parasite line. The parasites were cultivated at 37°C in an atmosphere of 5% oxygen, 5% carbon dioxide and 90% nitrogen at 5% hematocrit in RPMI 1640 medium, 0.5% Albumax II (Invitrogen), 0.25% sodium bicarbonate, and 0.1 mg/ml gentamicin. The parasites were synchronized using percoll/sorbitol gradient method in which infected RBCs were layered on a step gradient of 40/70 % percoll containing 6% sorbitol. The gradient was subsequently centrifuged at 12,000g for 20 minutes at room temperature. The late stage synchronized parasites were recovered from the interphase, washed twice with complete culture media and placed back in culture. The percentage of parasitemia was calculated by SYBRGreen I DNA stain (Life Technologies) using CytoFLEX (Beckman Coulter) Flow Cytometer.

### Bioinformatics analyses

The full-length sequence of histone H2A and its variants were obtained from different species based on sequence similarity. Multiple sequence alignment of PfH2A (PF3D7_0617800) and other histone H2A variants from different species were performed using CLUSTALW and further analyzed using ESPript3 program. A homology model of the PfH2A was built by comparative modeling using crystal structure of histone H2A (Protein Data Bank entry 1eqz, chain A) by using SWISS-MODEL server. The structure visualization of PfH2A 3D-model was performed using Pymol program.

### DNA damage of parasites by X-ray irradiation and H_2_O_2_

DNA damage in the parasites was performed by X-ray irradiation using a PXi precision X-ray irradiator set at 225 kV, 13.28 mA. In brief, 2 % ring stage NF54 parasites were exposed to different doses of X-ray irradiation (1000, 3000 and 6000 Rad). After irradiation parasites were either collected immediately (i.e 15 minute after irradiation) or put back in culture with fresh media for further analysis at different time points as mentioned elsewhere. The level of DNA damage was measured by In Situ DNA Fragmentation (TUNEL) Assay and phosphorylated H2A by western blot as described. To check the hydrogen peroxide (H_2_O_2_) mediated DNA damage, ring stage infected RBCs (~2 %) were treated with different concentrations of H_2_O_2_ (0-10, 50, 100 and 400 μM) for 1 hour at 37°C. Parasites were collected from RBCs by saponin lysis and the level of DNA damage was measured by phosphorylated H2A following western blot as described.

### In Situ DNA Fragmentation (TUNEL) Assay

Tightly synchronized ring stages parasites (NF54) were fixed for 30 minutes in freshly prepared fixative (4% paraformaldehyde and 0.005 % glutaraldehyde). After fixation cells were rinsed three times with PBS and incubated with permeabilization solution (0.1% Triton X-100 in PBS) for 10 min on ice. The cells were washed twice with PBS, and one time with wash buffer supplied with TUNEL Assay Kit-BrdU Red (Abcam cat # ab6610). TUNELassay was performed as per manufacturer guidelines. Briefly, following washing 50 μl of TUNEL reaction mixture (DNA labeling solution) was added to each sample. The cells were incubated for 60 min at 37°C with intermittent shaking. Cells were then washed three times with rinse buffer (5 min each time) and re-suspended in 100 μl of antibody solution for 30 minutes at room temperature. Cells were then washed three times with PBS and mounted using Invitrogen™ Molecular Probes™ ProLong™ Gold Antifade reagent with DAPI,and imaged using Nikon Eclipse Ti-E microscope equipped with a CoolSNAPMyo CCD camera..

### Western immunoblotting

Infected RBCs were lysed with saponin and parasites were pelleted down by centrifugation. The parasite pellet was subsequently washed twice with PBS and lysed in 2x Laemmli sample buffer. The protein lysates were centrifuged and the supernatants were subjected to SDS-PAGE (gradient 4-20%, Bio-Rad) and electroblotted to a nitrocellulose membrane. Immunodetection was carried out by using rabbit anti-γ-H2A.X (S^P^Q) primary antibody (generated using peptide containing the S^P^Q motif, Cell signaling cat # 9718S, 1:1000), anti-H2A antibody (Abcam cat# ab88770, 1:1000) and rabbit polyclonal anti-aldolase antibody (1:3000) [27]. The secondary antibodies used were antibodies conjugated to Horseradish Peroxidase (HRP), goat anti-rabbit (Jackson Immuno Research Laboratories, 1:10000). The immunoblots were developed in EZ/ECL solution (Israel Biological Industries).

### Immunofluorescence assay

Immunofluorescence assay (IFA) was performed as described previously with minor modifications[28]. In brief, iRBCs were washed twice with PBS and re-suspended in a freshly prepared fixative solution (4% Paraformaldehyde (EMS) and 0.0075% glutaraldehyde (EMS) in PBS) for 30 minutes at room temperature. Following fixation iRBCs were permeabilized with 0.1% Triton-X 100 (Sigma) in PBS, and then blocked with 3% BSA (Sigma) in PBS. Cells were then incubated with primary rabbit anti-γ-H2A.X (S^P^Q) (Cell signaling, cat # 9718S, 1:300) and anti H2A (Abcam cat# ab8870, 1:100) antibodies for 1.5 h at room temperature and washed three times in PBS. Following this, cells were incubated with Alexa Fluor 488 goat anti-rabbit (Life Technologies, 1:500) or Alexa Fluor 568 goat anti-rabbit (Life Technologies, 1:500) antibodies for 1h at room temperature. Cells were washed three times in PBS and laid on “PTFE” printed slides (EMS) and mounted in ProLong Gold antifade reagent with DAPI (Molecular Probes). Fluorescent images were obtained using a Plan Apo λ 100x oil NA=1.5 WD=130µm lens on a Nikon Eclipse Ti-E microscope equipped with a CoolSNAPMyo CCD camera. Images were processed using the NIS-Elements AR (4.40 version) software.

### Stochastic Optical Reconstruction Microscopy (STORM) imaging and analysis

STROM imaging was performed as described recently [29] using anti-H2A (Abcam cat# ab88770, 1:150) and rabbit anti-γ-H2A.X (S^P^Q) (Cell signaling, cat # 9718S, 1:300) as primary antibodies. Alexa Fluor 594 goat anti-rabbit (Life Technologies, 1:500) was used as a secondary antibody. Parasite nuclei were labeled with YOYO-1 (1:300, life technologies) for orientation and were not subjected to STORM. STORM was performed by a Nikon Eclipse Ti-E microscope with a CFI Apo TIRF × 100 DIC N2 oil objective (NA 1.49, WD 0.12 mm) as described. For each acquisition, 10000 frames were recorded onto a 256 × 256 pixel region (pixel size 160nm) of an Andor iXon-897 EMCCD camera. Super-resolution images were reconstructed from a series of the least 5000 images per channel using the N-STORM analysis module, version 1.1.21 of NIS Elements AR v. 4.40 (Laboratory imaging s.r.o.).

### Total Histone Extraction

Total histones were extracted using an acid extraction method as described previously with minor modifications [30]. All steps were performed at 4°C in buffers containing protease and phosphatase inhibitors to protect the enzymatic interference with PTMs. In brief, 200 ml of parasite cultures (~10% parasitemia) were saponin lysed and washed with PBS containing protease and phosphatase inhibitors. To prepare the intact nuclei, the cell pellet was re-suspended in 1 ml lysis buffer (20 mM HEPES pH 7.8, 10 mM KCl, 1mM EDTA, 1% Triton X-100 and 1mM DTT) and incubated for 30 min on rotator at 4 °C. Following cell lysis, the intact nuclei were washed and pelleted by centrifugation at 10,000g, for 10 min at 4 °C. The nuclei were re-suspended in 400 μl 0.4 N H_2_SO_4_ or 0.25 N HCl. The nuclei were incubated on a rotator overnight and supernatant containing the acid soluble histone fraction was collected after centrifugation at 16,000g for 10 min.

### Immunoprecipitation

Immunoprecipitation of PfH2A was performed as described (23) with slight modification. In brief, 200 ml of parasite cultures (∼10% parasitemia) were saponin lysed and washed with PBS containing protease and phosphatase inhibitors. Subsequently, the parasite pellet was dissolved in chilled lysis buffer containing 50 mM Tris/HCl pH 7.5, 150 mM NaCl, 1 mM EDTA, 0.01% SDS and 1% NP40 supplemented with protease and phosphatase inhibitors (Roche) and sonicated for 4-8 cycles of 10-15 sec at 45% output using Hielscher UP200S sonicator. The sonicated pellet was incubated for 30 minutes on ice. The lysate was purified by a few rounds of centrifugations at 10000g for 10 min. and incubated with primary antibody (anti H2A antibody (Abcam cat# ab88770) for 10-12 h at 4°C with continuous swirling. The supernatant was further incubated for 4–6 h with Protein A/G agarose beads (Pierce) at 4°C and beads were pelleted by centrifugation at 4°C. Beads were then washed with ice chilled washing buffer. Immunoprecipitated proteins were eluted with SDS Lamelli buffer and used for detection by SDS–PAGE and western blot analysis.

### Mass spectrophotometry (LC-MS/MS Analysis)

To identify the phosphorylated serine of PfH2A, the extracted histones were digested by trypsin and analyzed by LC-MS/MS on Q Exactive plus (Thermo). The peptides were identified by Discoverer software version 1.4 against the Plasmodium NCBI-NR database and against decoy databases (to determine the false discovery rate (FDR) using the sequest and mascot search engines. Semi quantitation was done by calculating the peak area of each peptide. The area of the protein is the average of the three most intense peptides from each protein. The results were filtered for proteins identified with at least 2 peptides with 1% FDR.

### KU-55933 inhibition assay

To check the effect of ATM kinase inhibitor (KU-55933) on intra erythrocytic parasite growth, and to calculate the IC50 (50% inhibitory concentration), a SYBR Green I based parasite growth assay was performed. KU-55933 was dissolved in DMSO to make a stock concentration of 10 mM and stored at (−20°C). For growth assay, sorbitol synchronized ring stage iRBCs were seeded in a 96 well plate at a final parasitemia of approximately 0.2 % and a hematocrit of 5 %. ATM kinase inhibitor (KU-55933) stock solution was diluted to 2X final concentration in complete RPMI medium and added to the prepared parasites at a volumetric ratio of 1:1. Each concentration of inhibitor was plated in triplicate and each well contained a total volume of 200 μl, with or without an appropriate concentration of KU-55933 (0-50 μM). In order to check the effects of DMSO, 1 μl of DMSO was added to all control wells. The plate was kept at 37°C in modular incubators that was gassed every 24 hours with 5% oxygen, 5% carbon dioxide and 90% nitrogen. After 72 hours, the media was discarded, and cells were washed twice with PBS. Parasite growth was determined by counting proportions of infected cells by SYBR Green I DNA stain (Life Technologies) using CytoFLEX (Beckman Coulter) Flow Cytometer. IC_50_ value was calculated by plotting % survival vs log inhibitor concentration using GraphPad Prism 6.0. Values were normalized and then curve fitted by non-linear regression. To check the effect of ATM kinase inhibitor (KU-55933) on histone PfH2A phosphorylation, Percoll/sorbitol synchronized late stage parasite were incubated with or without the inhibitor (20μM). After 24 hours, parasites were subjected to different dosages of X-ray irradiation (1000, 3000 Rad). After irradiation parasites were collected immediately (i.e 15 minute after irradiation) and subjected to western immunoblotting to check the level of phosphorylated H2A as described.

## Acknowledgments

This work was supported partially by the Israeli Academy for Science, Israel Science Foundation (ISF) Grant 1523/18 and in part by European Research Council (erc.europa.eu) Consolidator Grant 615412 and Ministry of Science and Technology Grant 89290 (to R.D.). RD is also supported by the Dr. Louis M. Leland and Ruth M. Leland Chair in Infectious Diseases. BS and MG were supported by the PBC Fellowship Program for Outstanding Post-Doctoral Researchers from China and India. VM is supported by the Minerva Stiftung for PhD students.

---

**Figure S1:**
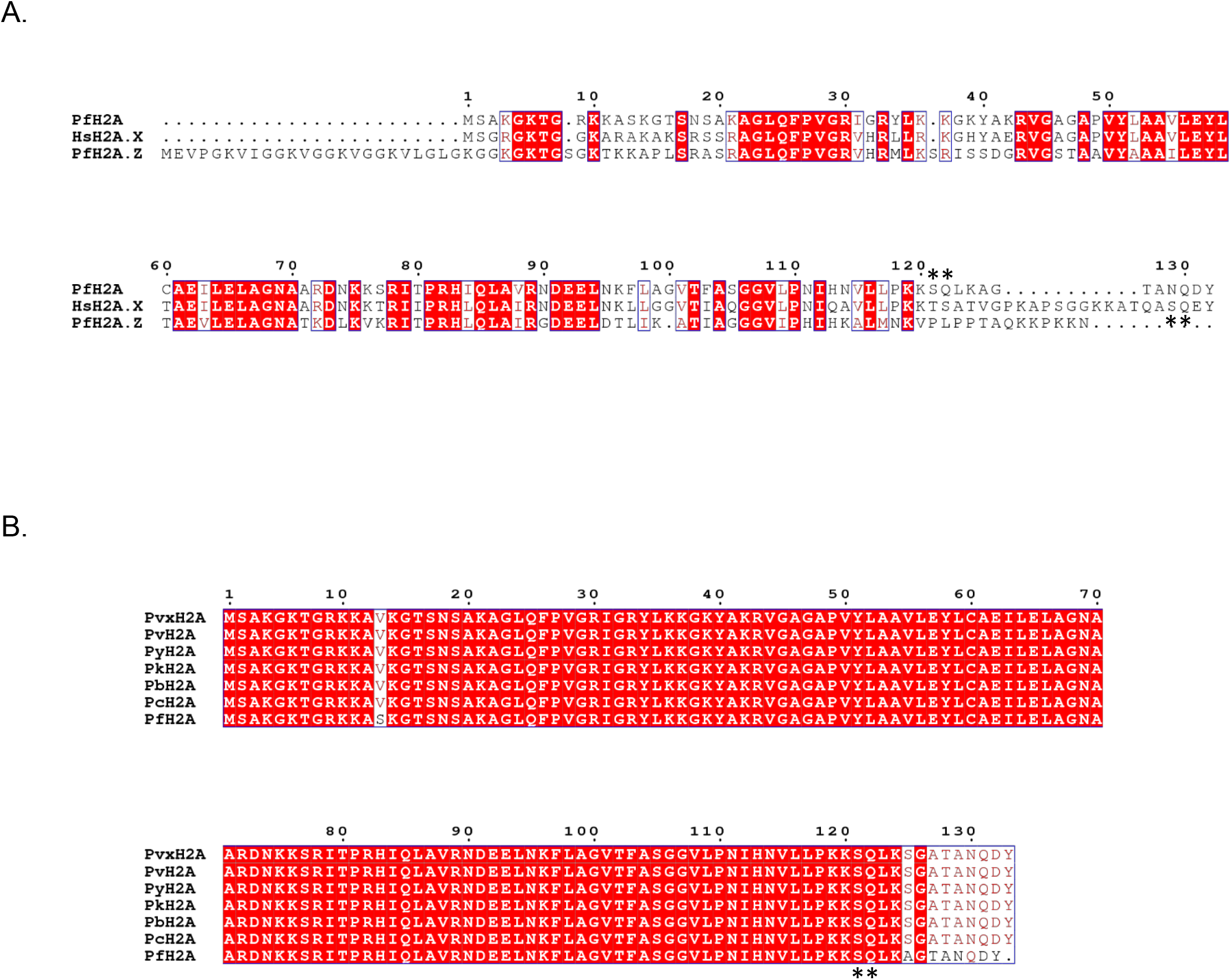
**(A)**. Multiple sequence alignment of amino acid sequences of histone H2A variants of *P. falciparum* and the human H2A.X, indicating that PfH2A.Z does not contain an SQ motif in its C’ tail. **(B)**. Multiple sequence alignment of amino acid sequences of histone H2A from different *Plasmodium* spp., indicating that the SQ motif in their C’ tail is highly conserved. The similar and identical amino acids are boxed and marked with a red background respectively. The conserved C-terminal SQ motif is marked with an asterisk. The species names and corresponding accession number are as follows: PfH2A; *Plasmodium falciparum* histone H2A (PF3D7_0617800), PfH2AZ; *Plasmodium falciparum* histone H2AZ (PF3D7_0320900) HsH2AX; *Homo sapiens* Histone H2A (NP_002096.1), PvxH2A; *Plasmodium vivax* histone H2A (PVX_114015), PvH2A; *Plasmodium vinckei* histone H2A (YYE_02539), PyH2A;, *Plasmodium yoelli* histone H2A (PY05076), PkH2A; *Plasmodium knowlesi* histone H2A (PKNH_1132600), PcH2A; *Plasmodium chabaudi* histone H2A (PCHAS_1116500), and PbH2A, *Plasmodium berghei* histone H2A(PBANKA_1117000), respectively.

**Figure S2:**
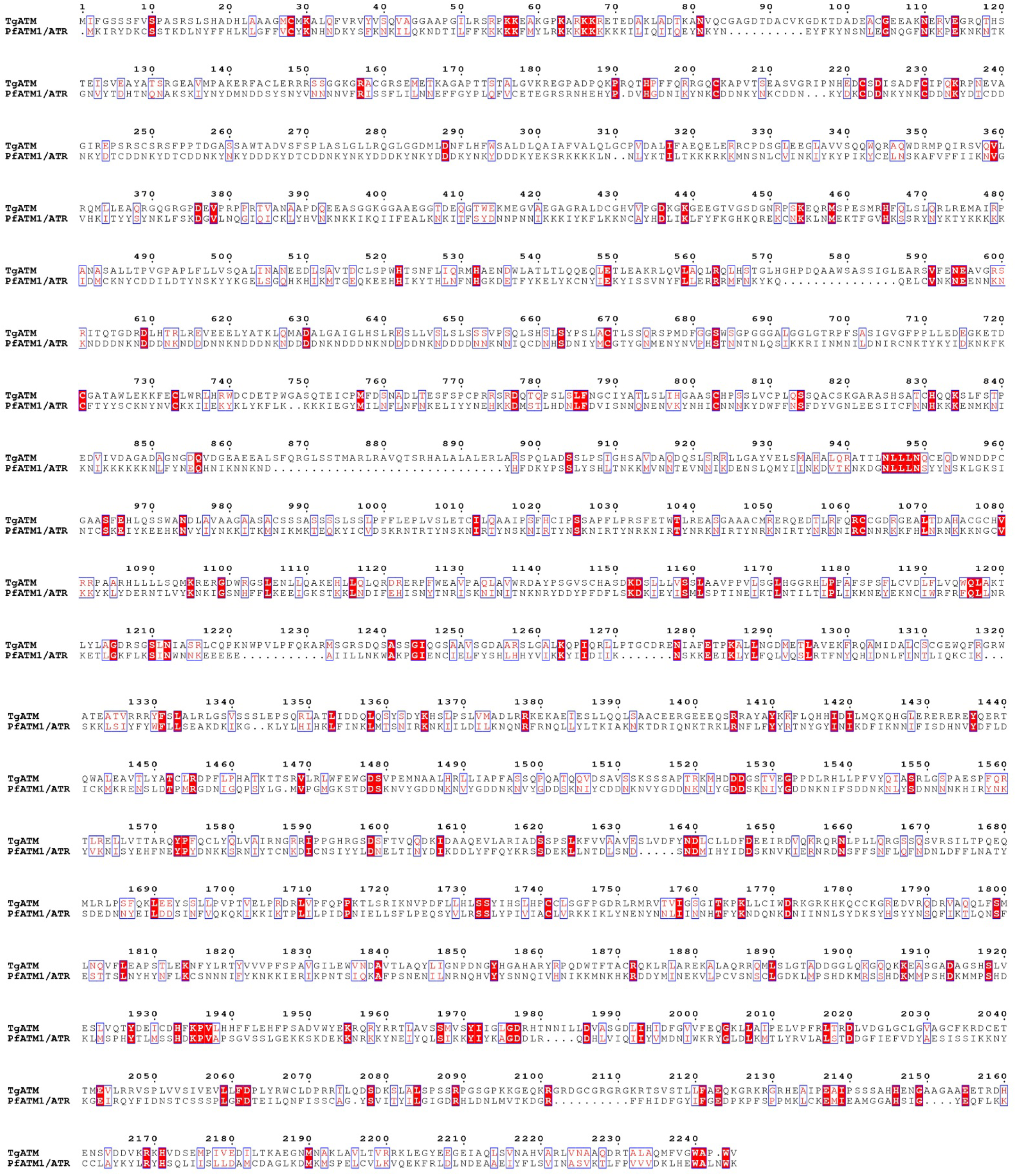
Multiple sequence alignment of amino acid sequences of putative ATM kinase from Toxoplasma and Plasmodium. The species names and corresponding uniport accession number are as follows: PfATM1/ATR; *Plasmodium falciparum* PF3D7_0515300, TgATM; *Toxoplasma gondii* TGME49_248530. The similar and identical amino acids are boxed and marked with a red background respectively.

**Figure S3.**
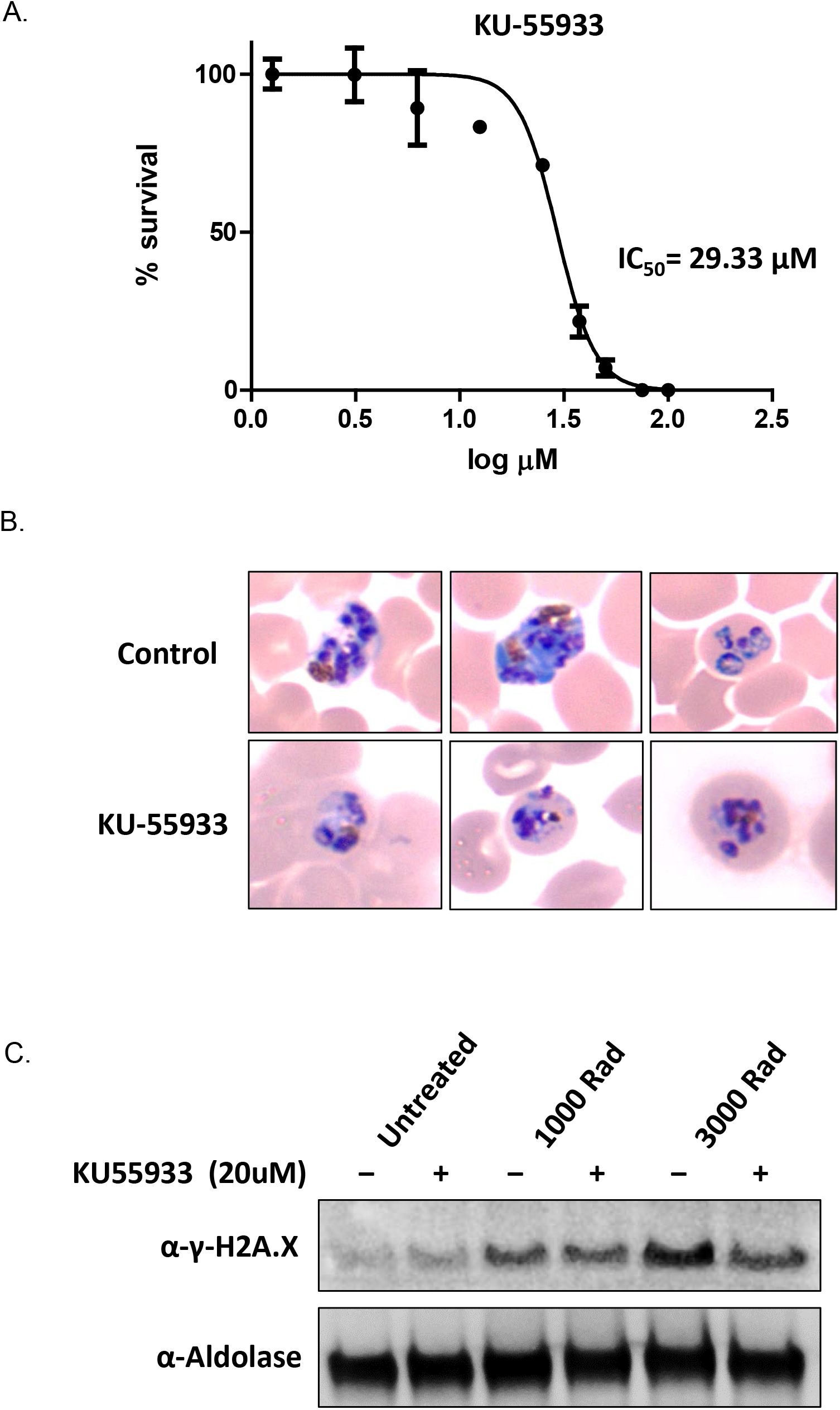
Effect of ATM kinase inhibitor KU-55933 on *Plasmodium falciparum* growth and PfH2A phosphorylation. **(A)**. *P. falciparum* iRBCs were treated with different doses of KU-55933 inhibitor (0-50 μM) for 72 h. After that, the iRBCs were washed with PBS, and the parasitemia was measured using SYBR Green. The IC50 value was calculated by plotting % survival vs log inhibitor concentration, normalized and curve fitted by non-linear regression. Data presented are mean of three biological replicates ±SD. The graph is representative of three independent experiments with similar results. **(B)**. Giemsa staining of parasites treated with KU-55933 inhibitor (25μM). **(C)**. *P. falciparum* iRBCs (Schizonts stage) were grown in the presence and absence of KU-55933 inhibitor (20μM) for 24 hours prior to exposure of iRBCs (ring stage) to different doses of X-ray radiation (i.e. 1000 and 3000 Rad or none). The levels of PfH2A phosphorylation with or without the inhibitor was measured by WB using anti-γ-H2A.X (S^P^Q) antibody and indicated a reduction when parasites were irradiated by 3000 Rad.

